# Bone morphogenetic protein 4 gene therapy in mice inhibits myeloma tumor growth, but has a negative impact on bone

**DOI:** 10.1101/575159

**Authors:** Marita Westhrin, Toril Holien, Muhammad Zahoor, Siv Helen Moen, Glenn Buene, Berit Størdal, Hanne Hella, Huipin Yuan, Joost D de Bruijn, Anton Martens, Richard WJ Groen, Fatima Bosch, Ulf Smith, Anne-Marit Sponaas, Anders Sundan, Therese Standal

**Author notes:** MW and TH contributed equally to this work. Correspondence: Toril Holien.

## Abstract

Multiple myeloma is characterized by accumulation of malignant plasma cells in the bone marrow. Most patients suffer from an osteolytic bone disease, caused by increased bone degradation and reduced bone formation. Bone morphogenetic protein 4 (BMP4) is important for both pre- and postnatal bone formation and induces growth arrest and apoptosis of myeloma cells. BMP4-treatment of myeloma patients could have the potential to reduce tumor growth and restore bone formation. We therefore explored BMP4 gene therapy in a human-mouse model of multiple myeloma where humanized bone scaffolds were implanted subcutaneously in RAG2^−/−^ γC^−/−^mice. Mice were treated with adeno-associated virus serotype 8 BMP4 vectors (AAV8-BMP4) to express BMP4 in the liver. When mature BMP4 was detectable in the circulation, myeloma cells were injected into the scaffolds and tumor growth was examined by weekly imaging. Strikingly, the tumor burden was reduced in AAV8-BMP4 mice compared with the AAV8-CTRL mice, suggesting that increased circulating BMP4 reduced tumor growth. BMP4-treatment also prevented bone loss in the scaffolds, most likely due to reduced tumor load. To delineate the effects of BMP4 overexpression on bone per se, without direct influence from cancer cells, we examined the unaffected, non-myeloma femurs by μCT. Surprisingly, the AAV8-BMP4 mice had significantly reduced trabecular bone volume, trabecular numbers, as well as significantly increased trabecular separation compared with the AAV8-CTRL mice. There was no difference in cortical bone parameters between the two groups. Taken together, BMP4 gene therapy inhibited myeloma tumor growth, but also reduced the amount of trabecular bone in mice. Our data suggest that care should be taken when considering using BMP4 as a therapeutic agent.

## Introduction

Multiple myeloma is a hematological cancer caused by accumulation of malignant plasma cells in the bone marrow (1). Nearly all patients suffer from a severe osteolytic bone disease, causing pain and fractures (2). The bone disease is caused by increased osteoclast activity and a lack of bone repair due to too few and dysfunctional osteoblasts (2). Currently, bisphosphonates are the most common drugs used to treat the bone disease, and there is a lack of treatment options that can promote bone formation. New, efficient drugs to treat myeloma have been developed the last decades, including immunomodulatory agents, proteasome inhibitors, histone deacetylase inhibitors, and monoclonal antibodies (3). Nevertheless, multiple myeloma remains an incurable disease (4).

Bone morphogenetic proteins (BMPs) is a large subgroup of ligands in the transforming growth factor (TGF)-β family (5). *In vitro*, several BMPs induce growth arrest and apoptosis in multiple myeloma cell lines as well as in primary myeloma cells from patients (6–11). BMP-signaling is also important for both pre- and postnatal bone formation.(12) For example, combined deletion of BMP2 and BMP4 in mesenchymal stem cells (MSC) leads to severely impaired osteogenesis in mice (13) and inhibiting BMP-signaling reduces osteoblast differentiation in mouse and human cells (14–16). On the other hand, two separate studies found increased bone mass and bone strength in mice treated with soluble BMPR1A-Fc fusion protein (17,18). BMPR1A-Fc has high affinity to BMP2 and BMP4 and acts as a decoy receptor that inhibits their binding to receptors on the surface of cells. Thus, the effects of a given BMP in the context of multiple myeloma is not entirely clear. In this study we wanted to clarify if BMP4 could have therapeutic potential in multiple myeloma patients, by preventing tumor growth and restoring bone homeostasis. We therefore evaluated the effects of AAV-based BMP4 gene therapy in a human-mouse scaffold model of multiple myeloma.

## Materials and Methods

### Cell culture and reagents

The human myeloma cell line KJON (19) was cultured in RPMI with 5 % heat inactivated human serum (HS) and 2 ng/mL interleukin (IL)-6 (Gibco, Thermo Fisher Scientific, Waltham, MA, USA). *In vitro* experiments with KJON cells were performed with 2 % HS in RPMI and IL-6 (1 ng/mL). The mouse myeloma cell line NS0 was generously provided by Dr Z. Eshhar (Weizmann Institute of Science, Israel), and the cells were grown in 10 % FCS in RPMI. All cells were cultured at 37 °C in a humidified atmosphere containing 5 % CO_2_ and were tested for mycoplasma every three months. Recombinant murine (rm) BMP4 (Cat# 5020-BP), recombinant human (rh) BMP4 (Cat# 314-BP), rhM-CSF (Cat# 216-MC), neutralizing BMP4 antibody (Cat# MAB50201), and rat isotype control (MAB006) were from R&D Systems (Bio-Techne, Abingdon, UK).

### Generation of iRFP-labeled KJON myeloma cells

The iRFP sequence was amplified from piRFP (gift from Vladislav Verkhusha, Addgene plasmid# 31857; http://n2t.net/addgene:31857; RRID:Addgene_31857)(20). with PCR primers with overhangs containing restriction sites for SpeI and NotI (Sigma). pLVX-EF1α-IRES-ZsGreen1 (Cat# 631982, Clontech, Takara Bio USA, CA, USA) was cut with SpeI and NotI restriction enzymes, treated with FastAP, and ligated with the iRFP PCR product, using T4 DNA ligase (all Fermentas, Thermo Fisher Scientific). The resulting plasmid, pLVX-EF1α-iRFP-IRES-ZsGreen1, was used together with TransLenti Viral Packaging Mix (Open Biosystems) and Genejuice (Novagen, Merck Life Science AS, Oslo, Norway) to transfect 293T packaging cells (Open Biosystems, Thermo Fisher Scientific). Supernatants containing lentivirus were used to transduce the KJON myeloma cell line. Positively transduced cells expressed both iRFP and ZsGreen1 fluorescent proteins and were sorted on a FACSAria™ Fusion flow cytometer (BD Biosciences, San Jose, CA, USA) to obtain a pure population of iRFP-positive cells for *in vivo* studies.

### Cell viability *in vitro*

Cell viability was measured using the Cell Titer-Glo assay (Promega, Madison, WI, USA), that measures the ATP content in wells. Cell lines were seeded in 96 well optical plates (10^4^ cells/well) and treated as indicated in the figure legends. Cell Titer-Glo reagent was added following the manufacturer’s instructions and luminescence was measured using Victor 1420 multilabel counter (PerkinElmer Inc., Waltham, MA, USA). To distinguish between effects on cell division and cell death, we measured apoptosis by annexin V labelling. In brief, cells were seeded in 96 well plates (5 x 10^4^ cells/well) and treated as indicated in the figure legends. The cells were stained using Apotest FITC kit (Nexins Research, Kattendijke, The Netherlands). Then, cells were incubated with annexin V FITC (0.2 μg/mL) on ice for one hour. Propidium iodide (PI) (1.4 μg/mL) was added five minutes before cells were analyzed using an LSRII flow cytometer (BD Biosciences, San Jose, CA, USA). Cells negative for both annexin-V and PI staining were considered viable.

### RAG2^−/−^ γC^−/−^

We used RAG2^−/−^ γC^−/−^ BALB/c female mice as described previously (21). The mice lack B, T and NK cell immunity and were kept in specific pathogen free (SPF) unit. Here the mice were housed in IVC-cages, with free access to bedding material, nesting material and enrichment objects. Mice were given sterile food (RM1 #801002, Special Diets Services, Essex, UK,) and water *ad libitum*, and were caged in groups of 3-5 mice. Mice were maintained at a room temperature of 21-22 °C and 55 % humidity with a 12 hr light/dark cycles including 1 hr dusk/dawn. All mice were of approximately the same age at the beginning of experiments.

### Human-mouse scaffold myeloma model and imaging

We used a modified version of a previously described xenograft mouse model with a humanized bone environment (21). Briefly, human bone marrow-derived mesenchymal stromal cells (hMSC) from healthy donors were seeded on biphasic calcium phosphate (BCP) scaffolds and differentiated towards osteoblasts for 1 week *in vitro*. Then, four cell-containing scaffolds were inserted subcutaneously on the back of 20 12-week old RAG2^−/−^ γC^−/−^ female mice and left for further differentiation of the cells. After 8 weeks, mice were treated with recombinant adeno-associated virus (AAV) of serotype 8, AAV8-CTRL (n=10) or AAV8-BMP4 (n=10) (10^12^ viral particles in 100 μL), by tail vein injections. These viruses contain the human α1-antitrypsin (*hAAT1*) promoter to ensure transgene expression in the liver. The AAVs were produced and purified as described previously.(22) After another 2 weeks, the mice were whole-body irradiated using a dose of 2 Gy photons on the day before injection of 10^6^ KJON cells into three of the scaffolds. The fourth scaffold was used as a non-tumor cell control. An overview of the experimental set-up is shown in Figure 1. Treatment with AAV is regarded relatively safe,(23) and in line with this, AAV8 treatment did not cause any adverse events in mice. However, one mouse in the AAV8-BMP4 group passed away for unknown reason. Tumor load (iRFP intensity) was measured weekly using Pearl Imager and analyzed with the accompanying Image Studio software (LI-COR Biosciences, Lincoln, NE, USA). At week 6 post myeloma cell injection, the mice were sacrificed, and serum, organs and bones were harvested. All mice were euthanized before tumors reached 1 cm^3^ (-scaffold) or when the tumor affected animal well-being. Animal handling and procedures were approved by the Norwegian food safety authority (FOTS7692). The experiment was not blinded.

**Figure 1.**
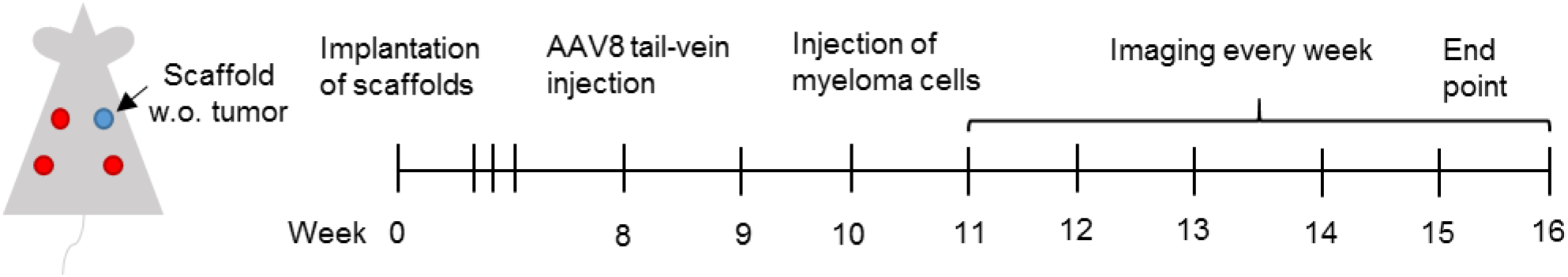
Experimental set-up. Overview of experimental set up. In brief, 14 weeks old RAG2^−/−^ γC^−/−^ mice were implanted with calcium phosphate scaffolds containing human MSCs. AAV8-BMP4 or AAV8-CTRL were administered 8 weeks post scaffold implantation by tail-vein injection (10^12^ viral particles/100 μL saline/mouse). 10 weeks post implantation 10^6^ fluorescently labeled KJON myeloma cells were injected into 3 out of 4 scaffolds. Empty scaffold refers to scaffold with MSCs, but without tumor cells.

### Western blotting

Mouse liver cells were prepared by cutting the liver in smaller pieces using a scalpel, before they were dissociated using gentleMACS™ Dissociator (Miltenyi Biotech, Bergisch Gladbach, Germany). Pelleted cells were lysed in lysis buffer (50 mM Tris-HCl (pH 7.5), 1 % IGEPAL CA-630 (Sigma-Aldrich), 150 mM NaCl, 10 % glycerol, 1 mM Na_3_VO_4_, 50 mM NaF and protease inhibitor cocktail (Roche, Basel, Switzerland)). For serum samples, 5 uL of serum was used per well. The samples were denatured in NuPage LDS sample buffer (Invitrogen, Thermo Fisher Scientific) supplemented with 25 mM dithiothreitol (DTT) for 10 min at 70°C before they were separated on 4-12 % Bis-Tris polyacrylamide gels with MES buffer (Invitrogen), and transferred to a nitrocellulose membrane using the iBlot Dry Blotting System (Invitrogen). The membrane was blocked using nonfat dry milk (5 %) diluted in Tris-buffered saline with 0.01 % Tween 20 (TBS-T). The primary antibodies were: Mouse anti-BMP4 (Cat# ab93939, RRID:AB_10562295) and mouse anti-GAPDH (Cat# ab8245, RRID:AB_2107448) (Abcam, Cambridge, UK). Blots were incubated with horseradish peroxidase (HRP) conjugated secondary antibodies (DAKO Cytomation, Glostrup, Denmark) and developed with SuperSignal West Femto Maximum Sensitivity Substrate (Thermo Fisher Scientific). Images were obtained with Odyssey FC and analyzed using Image Studio Software (LI-COR).

### Real-time RT-PCR

Liver cells were dissociated as described in the western blotting section and mRNA was isolated from the cells. Femurs from all mice were harvested, flushed and kept in liquid nitrogen until further processing. The femurs were then homogenized using Metal Bead Lysing Matrix (MP Biomedicals, LLC, OH, USA) and Trizol (Thermo Fisher Scientific). Samples of mRNA were reversely transcribed and RT-PCR analysis was performed using TaqMan Gene Expression Arrays (Applied Biosystems, Thermo Fisher Scienctific) as described previously.(24) The primers are listed in Supplementary Table 1. Genes with a Ct value ≥ 36 were considered as not detected. StepOne Software v2.1 (Applied Biosystems) was used to analyze the samples and the comparative Ct method was used to estimate relative changes in gene expression using *Gapdh* as housekeeping gene.

### Osteoclast differentiation

Peripheral blood mononuclear cells (PBMCs) were isolated from healthy donors using Lymphoprep (Axis-Shield, Oslo, Norway). CD14^+^ cells were further purified using magnetic beads (Miltenyi). Cells were seeded out in 96 well plates and cultured in aMEM with 10 % heat inactivated HS and M-CSF (30 ng/mL) for 2 days. At this point rhBMP4 (20-200 ng/mL) was added as indicated in the figure legend. When multinuclear cells were visible with light microscopy, cells were fixed and stained for TRAP using Acid Phosphatase, Leukocyte (TRAP) kit, (Merck KGaA, Darmstadt, Germany). TRAP positive cells with 3 or more nuclei were counted.

### Scaffold bone analysis

The scaffolds were harvested at end point and decalcified using Osteosoft (Merck). After approximately 4 weeks, when scaffolds were soft to the touch, they were embedded in paraffin, sectioned (3.5μm) and stained with hematoxylin and eosin. Images were acquired using Nikon Microscope ECLIPSE Ci-S. The amount of bone and total scaffold perimeter were quantified using NIS Elements (BR 4.00.00, Nikon).

### uCt analysis

Femurs were harvested and examined by *ex vivo* μCT using a μCt scanner (Skyscan 1176, Bruker, Kontich, Belgium) as described.(25) Images were acquired using the following settings: 18-μm voxel resolution, 0.5-mm aluminum filter, 50-kV voltage, and 500-μA current, 252 ms exposure time, rotation 0.5 degrees, frame averaging = 4. Images were reconstructed and analyzed using SkyScan software programs NRecon (version 1.6.9.4), DataViewer (version 1.4.4), and CT Analyzer (version 1.12.10.0). Femoral trabecular analysis region of interest (ROI) was determined by identifying the distal end of the femur and calculating 10 % of the total femur length toward the femora mid-shaft, where we then analyzed an ROI of 15 % of the total femur length. Analysis of bone structure was completed using adaptive thresholding (mean of minimum and maximum values) in CT Analyzer. Thresholds for analysis were determined manually based on grayscale values (0–255, where 0 = black and 255 = white) and were set as 36 to 255. Cortical analyses were performed 35 % above the distal end of the femur toward the femora mid-shaft, also with a 15 % ROI with the threshold values set as 80 to 255.

Images were generated using CtVox (version 3.3) (Skyscan 1.1.6.0).

### Statistical analysis

Statistical analysis was performed using GraphPad Prism 7.04. To compare two groups we used the unpaired, two-tailed t-test. To compare more than 2 groups we performed 1- or 2-way ANOVA with Dunn’s test or Bonferroni Post hoc test, respectively. Results were considered significant when p<0.05.

## Results

### BMP4 gene therapy reduces tumor growth *in vivo*

BMP4 induces apoptosis in approximately half of primary myeloma cell samples tested, as well as in a few myeloma cell lines (7,9). Here, we wanted to use a cell line which is both sensitive to BMP4 and relies on a supportive tumor microenvironment to grow *in vivo*. KJON is a slowly proliferating, IL-6 dependent cell line that is sensitive to BMP9 (10,19). BMP4 induces apoptosis in this cell line *in vitro* (Figure 2A, B). To investigate if BMP4 would also reduce *in vivo* tumor growth, we utilized an adeno-associated virus vector expressing BMP4 under the control of a liver-specific promoter (AAV8-BMP4) (22) in a human-mouse scaffold model of multiple myeloma (21). AAV8-BMP4 or control (AAV8-CTRL) viral particles were injected into the tail veins of mice that were implanted with scaffolds seeded with human bone marrow-derived MSCs8 weeks earlier. 2 weeks after virus injection, when recombinant BMP4 was detectable in blood plasma (data not shown), fluorescently labeled KJON myeloma cells were injected directly into 3 out of 4 scaffolds in each mouse.

**Figure 2.**
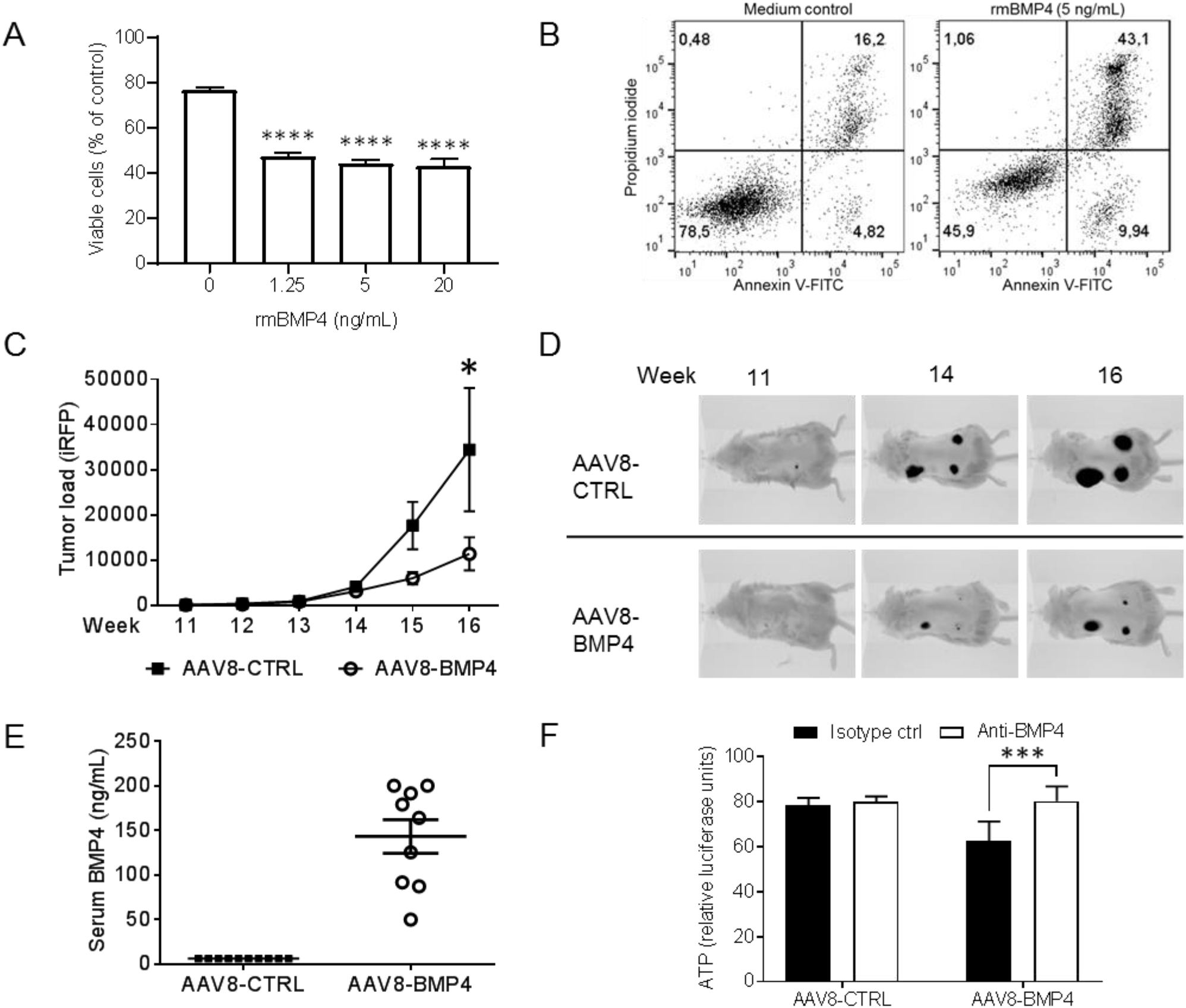
BMP4 inhibited myeloma cell growth *in vivo*. (A) The myeloma cell line KJON was treated with different doses of rmBMP-4. After 48 h, the numbers of viable cells were determined by labeling with annexin V-FITC and propidium iodide (PI). (B) The figure shows representative dot plots of the results presented in A. Cells in the lower left quadrant, which were negative for both annexin V and PI, were considered viable. (C) To estimate tumor burden, the amount of near-infrared fluorescent protein (iRFP) in each scaffold was measured weekly using the Pearl Imager System in AAV8-CTRL mice (n=30) and AAV8-BMP4 mice (n=27), p<0.01, 2-way ANOVA, Bonferroni post test. (D) Representative images of tumor burden in AAV8-CTRL (top) and AAV8-BMP4 (bottom) treated mice are shown. (E) Amount of BMP4 in the serum was estimated at end point by semi-quantitative western blotting. (F) Serum from AAV8-CTRL or AAV8-BMP4 mice, 4% final serum concentration, was added to cultures of NS0 murine myeloma cells and incubated for 48 h. Reduced cell proliferation, as measured by ATP-levels, was counteracted by a BMP4 neutralizing antibody, p<0.0005, Bonferroni post test.

Tumor growth was examined by weekly imaging, until 6 weeks after myeloma cell injection when the mice were euthanized. We found a significant reduction in tumor load in AAV8-BMP4 mice compared with the AAV8-CTRL mice (p<0.01, Figure 2 C, D). At this time point, serum levels of BMP4 were high (50-200 ng/mL) in the AAV8-BMP4 mice (Figure 2E). The BMP4 was biologically active, since addition of sera obtained from the AAV8-BMP4-treated mice to the murine myeloma cell line NS0, led to reduction in cell viability. Adding BMP4-neutralizing antibody restored viability (Figure 2F). In contrast, NS0 viability was not affected by sera obtained from AAV8-CTRL mice. Further supporting successfully transduction, we could detect the pro-form of BMP4 in liver lysates from AAV8-BMP4 mice, but not from AAV8-CTRL mice (Supplementary Figure 1A, B), and we also observed that AAV8-BMP4 treatment increased downstream BMP targets such as *Smad7* and *Id1* in the liver (Supplementary Figure 1 C, D). To examine if the BMP4 transgene would alter the expression of *endogenous* BMP4 or other BMPs, we analyzed the mRNA expression of mouse *Bmp4, Bmp2, Bmp6, Gdf2* (BMP9) in the liver. The expression of these did not change (Supplementary Figure 1 E-H). Taken together, these data demonstrated that AAV8-BMP4 gene therapy lead to the production of high amounts of circulating, biologically active BMP4 that reduced tumor growth.

Continuous drug exposure may generate acquired resistance in multiple myeloma (26). To examine if the tumor cells became resistant to BMP4 during the treatment, we isolated live cells from tumors from both AAV8-BMP4 and AAV8-CTRL mice. Cells from both groups were equally sensitive to BMP4 treatment *in vitro*, indicating that they did not acquire resistance to BMP4 during the experiment (Supplementary Figure 2).

### BMP4-treatment rescues bone loss in the scaffolds

Human bone is generated on the scaffolds by osteoblasts that differentiate from human MSCs seeded on the scaffolds before implantation (21). In this model, similar to what happens in multiple myeloma, bone formation in the scaffolds is impaired by the presence of myeloma cells (21). This was also the case here, as we found less bone in scaffolds with myeloma cells compared with the scaffolds without myeloma cells (Figure 3A, left, p<0.05). Interestingly, this difference was lost in the AAV8-BMP4-treated mice (Figure 3A, right panel). This could be explained by an increased availability of BMP4 in the AAV8-BMP4 treated mice, promoting osteoblast differentiation and bone formation in the humanized bone scaffolds. However, when we quantified amount of bone in scaffolds without tumors, we found no significant difference between AAV8-BMP4 treated mice compared with AAV8-CTRL mice (Figure 3A). Thus, it is likely that the beneficial effect on bone formation in the scaffolds is due to reduced tumor burden in AAV8-BMP4-treated mice.

**Figure 3.**
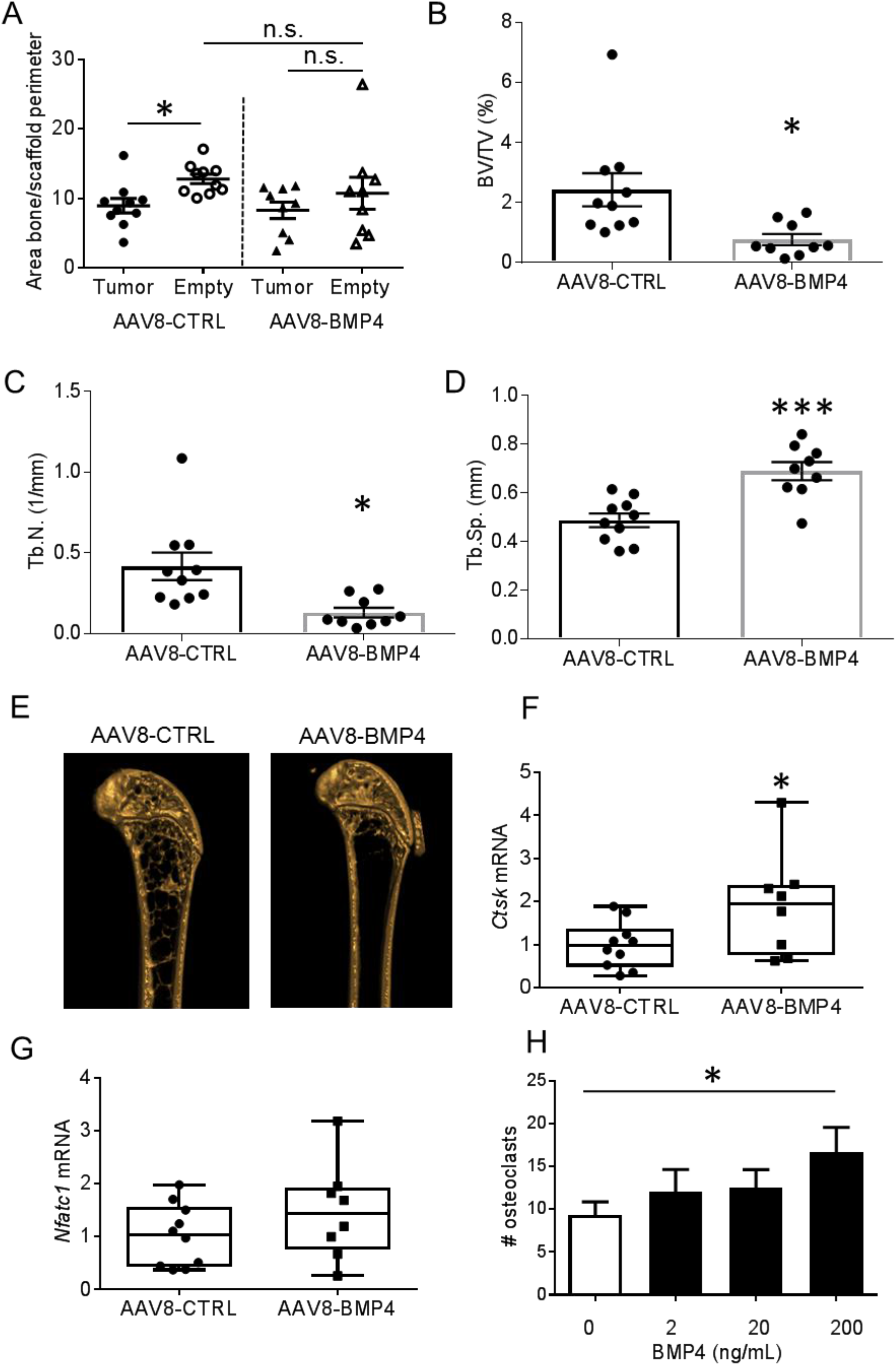
BMP4 effect on human and murine bone. (A) Amount of bone/scaffold perimeter is presented for both empty and tumor scaffold for AAV8-CTRL treated mice (n=10) and AAV8-BMP4 treated mice (n=9). The left femur from each mouse, AAV8-CTRL (n=10) and AAV8-BMP4 (n=9), was harvested and examined by *ex vivo* μCT. *; p<0.05, 1-way ANOVA, Dunn’s multiple comparisons test. (B) Trabecular volume as a proportion of tissue volume (BV/TV, %), (C) trabecular number (Tb. N, mm^−1^) and (D) trabecular separation (Tb.Sp, mm) was assessed. Error bars represent SEM. *; p<0.05, ***; p<0.005, two-tailed unpaired t-test. (E) Representative images for an AAV8-CTRL mouse and an AAV8-BMP4 mouse are shown. Femur cDNA from AAV8-CTRL mice (n=10) and AAV8-BMP4 mice (n=8) was used for comparative RT-PCR using TaqMan Assays for the osteoclast specific markers *Ctsk* (F) and *Nfatc1* (G). The relative gene expression was analyzed using the ΔΔCt method with *Gapdh* as housekeeping gene. *; p<0.05, two-tailed unpaired t-test. (H) Cells were differentiated with M-CSF (30 ng/mL) in the presence or absence of rmBMP4 as indicated. TRAP positive cells with more than 2 nuclei were counted as osteoclasts. Presented is the mean of three independent experiments and error bars represent SEM. *; p<0.05, 1-way ANOVA, Dunn’s multiple comparisons test.

### High circulating levels of BMP4 reduces mouse trabecular bone volume

The myeloma cells were confined to the scaffolds, as we could not detect plasma cells in the spleen or murine bone marrow when examined by imaging or flow cytometry (data not shown). We next aimed to investigate the effects of the elevated levels of circulating BMP4 on bone alone, without influence from the cancer cells. First, we established that the tumors did not affect the mouse bones by some soluble factor secreted from the tumor cells. This was done by comparing cortical and trabecular bone parameters in femurs obtained from un-implanted control mice (untreated RAG2^−/−^ γC^−/−^) with the AAV8-CTRL mice. As shown in Table 1 there was no difference in trabecular or cortical bone parameters between the groups of mice when examined by *ex vivo* μCT. Thus, the presence of tumor/scaffold had no apparent effect on mouse bone and the femurs in the AAV8-CTRL mice can be considered “naïve”.

**Table 1.** Trabecular bone parameters in untreated RAG2^−/−^ γC^−/−^ vs AAV8-BMP4 mice.

| | Mean ± SEM | | p-value (Unpaired t-test) |
| --- | --- | --- | --- |
| | AAV8-CTRL | RAG2<sup>-/-</sup> γC<sup>-/-</sup> | AAV8-CTRL vs RAG2<sup>-/-</sup> γC<sup>-/-</sup> |
| BV/TV (%) | 2.425±0.55 | 3.291±0.94 | 0.412 |
| Tb.Th (mm) | 0.05684±0.00 | 0.05575±0.00 | 0.5448 |
| Tb.Sp (mm) | 0.4873±0.03 | 0.4668±0.04 | 0.6882 |
| Tb.N/mm | 0.4166±0.09 | 0.5843±0.16 | 0.3302 |

We next examined the effect of BMP4 on mouse bone by comparing properties of femurs obtained from AAV8-BMP4 treated mice with the AAV8-CTRL treated mice. Surprisingly, the AAV8-BMP4 mice had significantly reduced trabecular bone volume (bone volume per tissue volume; % BV/TV, p=0.017), trabecular numbers (Tb.N/mm, p=0.016) as well as significantly increased trabecular separation (Tb.Sp (mm)) compared with the AAV8-CTRL mice (p=0.0005, Figure 3B-E). Thus, high levels of circulating BMP4 were detrimental for trabecular bone. In contrast, there were no differences in cortical bone parameters between the two groups (Supplementary Figure 3). Moreover, since RAG2^−/−^ γC^−/−^ mice are immunocompromised, we performed a small study using the same viral vectors in female immunocompetent C57BL6/N mice (n=4/5 group). Although the reduction in bone volume (% BV/TV) in AAV8-BMP4 compared to AAV8-CTRL mice was not significant, there was a significant increase in trabecular separation (Tb.Sp) and a reduction in trabecular numbers (Tb.N/mm) in AAV8-BMP4 treated mice compared to AAV8-CTRL mice, supporting that high levels of BMP4 may have a negative impact on trabecular bone also in immunocompetent mice (Supplementary Figure 4).

### Effects of BMP4 gene therapy on osteoblasts and osteoclasts

In an attempt to determine whether the BMP4-induced effects on bone were caused by increased bone resorption or decreased bone formation we measured the bone degradation marker C-terminal telopeptide of Type I collagen (CTX-1) and bone formation marker type I pro-collagen N-terminal pro-peptide (PINP) in mouse sera (Supplementary Figure 5). However, there were no significant differences between the groups. We also analyzed the murine femurs to examine if high BMP4 levels had altered osteoblast differentiation. Again, mRNA expression of osteocyte- and osteoblast-specific markers (Sclerostin (Sost), Dickkopf-related protein 1 (*Dkk1*), Runt-related transcription factor 2 (*Runx2*) and Osterix (*Sp7*)) did not differ between the groups (Supplementary Figure 6). Although this may suggest that osteoblast differentiation was not significantly affected, the variation in gene expression within groups were high, which makes it hard to conclude on this matter.

For the osteoclast-specific markers, Cathepsin K (*Ctsk*) and Nuclear factor of activated T-cells, cytoplasmic 1 (*Nfatc1*), we found a significant increase in *Ctsk* in the AAV8-BMP4 mice (p<0.05, Figure 3F, G). We therefore investigated if BMP4 had an osteoclast-promoting effect *in vitro*. Indeed, addition of recombinant human (rh) BMP4 to CD14^+^ osteoclast-precursors increased osteoclast differentiation (Ctrl vs rhBMP4 (200 ng/mL), p<0.05, Figure 3H). Taken together our results infer that BMP4 has a negative impact on bone, at least in part, by increasing osteoclast numbers.

## Discussion

In this study we wanted to explore BMP4 gene therapy as a potential treatment for multiple myeloma in a human-mouse model. We found that BMP4 gene therapy inhibited myeloma tumor growth, but surprisingly reduced trabecular bone in mice.

BMPs, like TGF-β, usually act as tumor suppressors (5). In multiple myeloma, several different BMPs inhibit growth *in vitro* (6–10). The abundance of different BMPs in the bone marrow is not known. It was shown that *BMP6* mRNA is expressed by both normal and malignant plasma cells, and that high levels of *BMP6* in myeloma cells was associated with a favorable prognosis in multiple myeloma patients (27). This suggests that BMPs can have anti-tumor effects in patients. We show here that BMP4 treatment inhibited tumor growth *in vivo*, in line with previous studies examining effects of BMP4 on tumor cell survival *in vitro*. Importantly, more than half of the patient-derived primary cells we tested *in vitro* were sensitive to BMP4, which implies that BMP4 could have beneficial effects in a large group of patients (7,9). Multiple myeloma is a very heterogeneous disease, and the malignant cells harbor different genetic aberrations (27). About 50 % of all myeloma patients have cancer cells that are hyperdiploid (27). Despite this high number of hyperdiploid cells, very few cell lines have been established with this genotype. Here, we used the hyperdiploid myeloma cell line, KJON, which has a relatively slow growth rate and relies on addition of recombinant IL-6 in the absence of a supporting microenvironment, thus resembling what takes place in patients (19). Such cells are usually not able to grow in a mouse bone marrow microenvironment (28), but in this model the human mesenchymal cells provide the support needed. The model also recapitulates the tumor-induced bone loss, which is a characteristic feature of myeloma. Thus, we here show that BMP4 gene therapy inhibited myeloma growth in a relevant *in vivo* model. Importantly, we found that reduction of tumor size by BMP4 gene therapy had a beneficial effect on bone, as it prevented loss of bone in tumor-affected bone, although it did not promote bone formation. However, it is important to note that the human-mouse scaffold model cannot be used to study tumor-induced bone loss in load-bearing bone.

The bone-inducing effect of BMPs has been known since the 1960s, and BMP2, BMP4-7 and BMP9 have all been appreciated for their osteogenic potential.(12,29) We were therefore surprised to find that trabecular bone was significantly reduced in AAV8-BMP4 treated mice compared with AAV8-CTRL mice, and that high levels of circulating BMP4 failed to promote bone formation in the scaffolds. To examine if this effect was somehow specific for RAG2^−/−^ γC^−/−^ BALB/c mice, we performed a small study in immunocompetent C57BL6/N mice using the same AAV8-BMP4 vector. Also in these mice, overexpression of BMP4 decreased the amount of trabecular bone, while cortical bone was unaffected. Our results are thus in contrast to previous studies demonstrating that BMPs, including BMP4, promote osteoblast differentiation and bone formation.(13–15,30). Further, in another study researchers found that injection of AAV2-BMP4 into the skeletal muscle of immunocompetent rats resulted in new bone induced by endochondral ossification already at week 3 (31). In contrast to our study, where we used AAV8-BMP4 to increase circulating levels of BMP4, it is likely that the AAV2-BMP4 mainly increased BMP4 locally, thus explaining the different outcomes. On the other hand, and in line with our data, BMP4 overexpression in bone caused severe osteopenia and increased osteoclast number in mice (32). The same study also found that overexpression of noggin, a BMP antagonist, had opposite effects. BMP2 and BMP4 have high affinity for BMPR1A, and treating mice with a decoy receptor for these BMPs (i.e. a BMPR1A Fc-fusion protein) also led to increased numbers of osteoblasts and reduced numbers of osteoclasts, resulting in higher bone mass in these mice (17,18). Supporting these data, conditional deletion of *Bmpr1a* in osteoblast-lineage cells has been shown to increase bone mass in mice (33,34). In conclusion, our results that BMP4 has a negative effect on trabecular bone in mice is supported by previous reports.

A weakness of our study is the lack of bone histomorphometric data. We can therefore only speculate on how BMP4 acted at the cellular level to reduce trabecular bone. Expression of mRNA in femurs flushed of bone marrow showed significantly higher expression of *Ctsk* in the AAV8-BMP4 treated mice, which may indicate that there were more osteoclasts in these bones. There was, however, a great variation in gene expression within groups and no differences in serum CTX-1 levels, so we could not make a conclusion on this matter. While BMPs in bone have mainly been studied in relation to effects on the osteoblast lineage, a few studies have earlier shown that BMP2, BMP4, BMP5, and BMP6 promote osteoclast differentiation *in vitro* and/or *in vivo*, supporting our findings (32,35,36). An indirect effect of BMP2 and BMP4 on osteoclasts was shown in a study where *Bmpr1a* was conditionally deleted in osteocytes leading to decreased RANKL expression in osteocytes and reduced osteoclast differentiation (33). The importance of osteoblastic BMP-signaling for RANKL expression was also shown in other studies (17,34). Together these studies and our data presented herein support that BMP-signaling impacts bone remodeling by influencing both osteoblast and osteoclast differentiation.

AAV mediated gene delivery methods are considered safe and have high gene delivery efficacy and are therefore promising tools for gene therapy (37). In adult mice, the same AAV8-BMP4 vector as used here was shown to increase insulin sensitivity and protected mice on a high-fat diet from obesity (22). In line with this study,(22) we found that while control mice had a weight gain of about 5 % in the 8 weeks from viral injections until culling, the weight of BMP4 treated mice remained unchanged during the course of the study. However, it is unlikely that such a small difference in body weight will lead to a dramatic effect on trabecular bone. Moreover, in the C57BL6/N immunocompetent mice there was no difference in body weight between the groups, and we could still observe a reduction in trabecular bone.

Taken together, BMP4 gene therapy inhibited myeloma tumor growth, but also reduced the amount of trabecular bone in mice. Care should therefore be taken when considering using BMP4 as a therapeutic agent. Whether other BMPs that are also potent inhibitors of multiple myeloma cell survival and proliferation will have similar impact on bone remains to be investigated.

## Authorship Contribution

MW and TH designed the study, performed experiments, analyzed data and wrote the paper; SHM and MZ assisted with mouse experiments; GB assisted with mouse experiments and analyzed data; BS and HH performed experiments; YH/JDdB provided scaffolds; AM/RWJG provided mice and designed the study; FB provided virus and designed the study; US/AS designed the study; TS designed the study, analyzed data and wrote the paper. All authors revised the manuscript before submission.

## Acknowledgements

This work was supported by funds from the Norwegian Cancer Society (#5793765 and #4500930), the Norwegian Research Council (#223255 and #193072), the Liaison Committee for education, research and innovation in Central Norway (90061001) and Ministerio de Economía y Competitividad (MINECO) (SAF2014-54866R), Spain. The authors would like to thank Kristine Misund for cloning the pLVX-EF1α-iRFP-IRES-ZsGreen1 plasmid. Parts of the *in vivo* work was provided by the Comparative Medicine Core Facility (CoMed), Norwegian University of Science and Technology (NTNU). CoMed is funded by the Faculty of Medicine at NTNU and Central Norway Regional Health Authority.

