## Supplemental data for "Bone morphogenetic protein 4 gene therapy in mice inhibits myeloma tumor growth, but has a negative impact on bone"

### Supplementary data

**Supplementary Materials and Methods**

**Supplementary Figure 1. BMP4 was expressed in liver of AAV8-BMP4 mice.**

**Supplementary Figure 2. KJON cells remained sensitive to BMP4 after the *in vivo* experiment.**

**Supplementary Figure 3. BMP4 effect on murine cortical bone.**

**Supplementary Figure 4. BMP4 effect on C57BL6/N murine bone.**

**Supplementary Figure 5. CTX-1 and PINP serum values**

**Supplementary Figure 6. Expression of osteoblast specific markers in femurs.**

**Supplementary Table 1. Taqman primers used.**

#### Supplementary Materials and Methods

### C57BL6/N

Female C57BL6/N were obtained from the Janvier laboratories (Le Genest-Saint-Isle, France). They were housed at our facilities with free access to bedding material, nesting material and enrichment objects. Food (RM1 #801002, Special Diets Services) and water was provided ad libitum. The mice were caged in groups of 5. All mice were at the same age (8 weeks) at the start of the experiment. At this point mice were treated with recombinant AAV8-CTRL (n=5) or AAV8-BMP4 (n=5) ($10^{12}$ viral particles in 100 µL), by tail vein injections. After 10 weeks the mice were euthanized and the femurs examined by µCT.

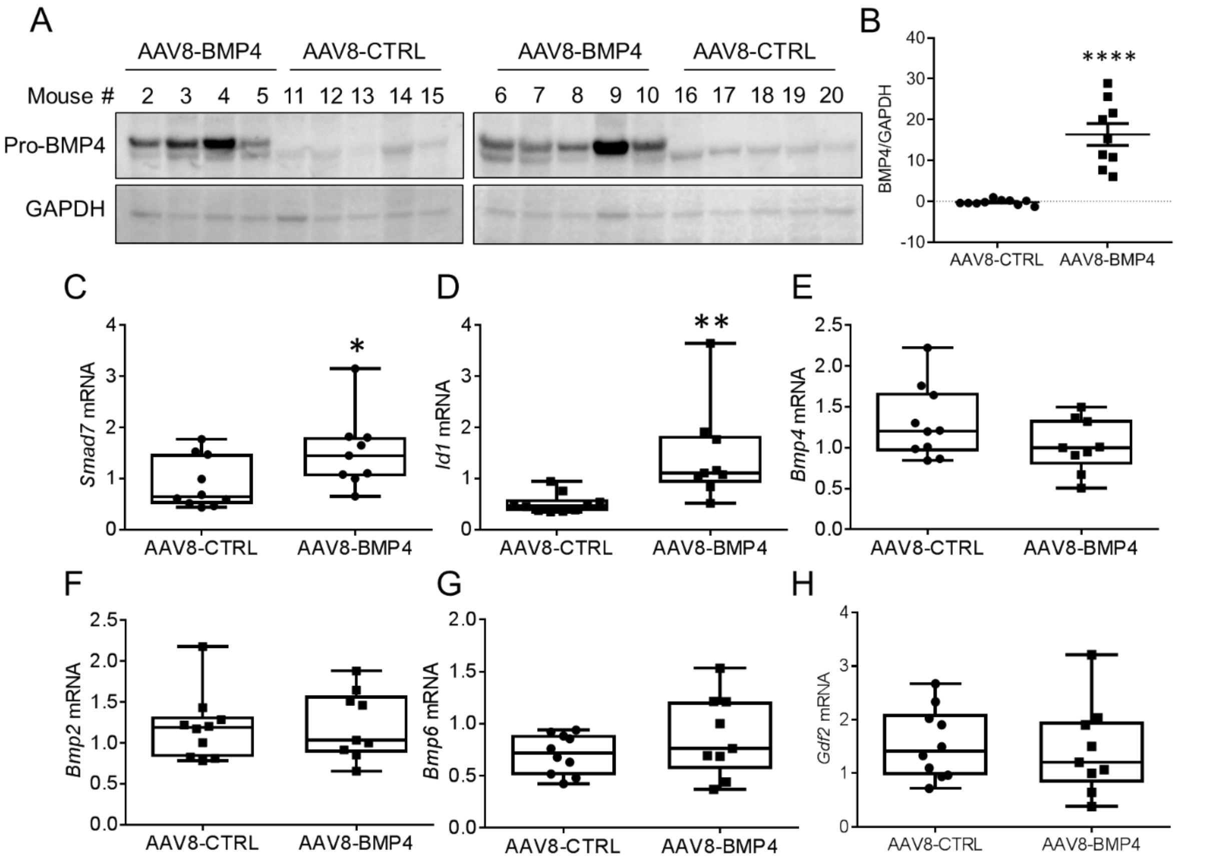


**Supplementary Figure 1. BMP4 was expressed in liver of AAV8-BMP4 mice.** A. Liver cells from mice were isolated using MACS dissociator (Miltenyi), lysed and used for immunoblotting with primary antibody targeting BMP4. A band at approximately 55 kDa was present in AAV8-BMP4 mice (n=9), but not in AAV8-CTRL mice (n=10), indicating the presence of pro-BMP4. B. Expression of pro-BMP4 relative to GAPDH was calculated and the values were compared. C-H. Other aliquots of dissociated liver cells were used for mRNA isolation and cDNA preparation using standard methods. RT-PCR analysis was performed using TaqMan Gene Expression Assays for *Smad7*, *Id1*, *Bmp4*, *Bmp2*, *Bmp6*, *Gdf2* (encoding BMP9), and *Gapdh*. Relative gene expression was analyzed using the ΔΔCt method with *Gapdh* as housekeeping gene. Of note, the virally expressed *Bmp4* is codon-optimized and thus the *Bmp4* TaqMan Assay used here only detects endogenous levels. Two-tailed, unpaired t-tests were used for statistical analysis, *; $p<0.05$, **; $p<0.01$, ****; $p<0.001$.

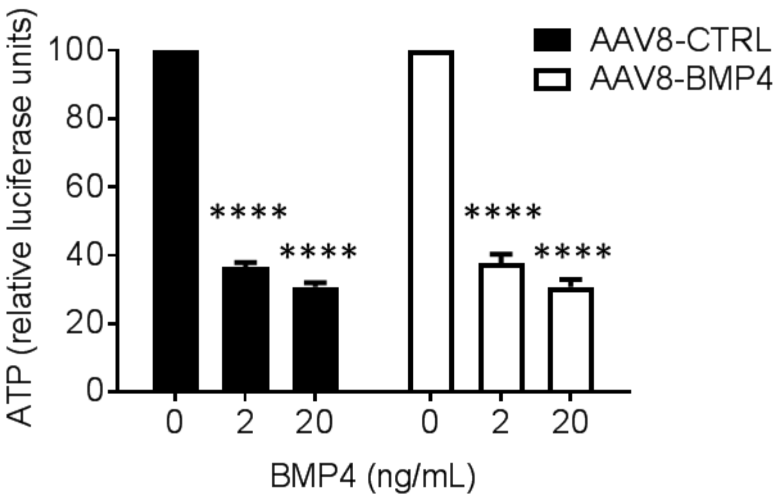


**Supplementary Figure 2. KJON cells remained sensitive to BMP4 after the *in vivo* experiment.** Tumor cells from AAV8-CTRL mice (n=5) and AAV8-BMP4 mice (n=5) were removed from the scaffolds and treated with recombinant murine (rm) BMP4 for three days. Sensitivity to BMP4 was evaluated using CellTiter Glo. Bars indicate normalized mean values compared with control (100%). There was no significant difference in ATP levels between medium control (BMP4 0 ng/mL) tumor cells from AAV8-CTRL mice and AAV8-BMP4 mice before normalization (unpaired two-tailed t-test, data not shown). Significance between the different BMP4-treated cells was calculated using two-way ANOVA, Bonferroni multiple comparisons test, ****; $p<0.001$. There was no significant difference in BMP4-effect between tumor cells from AAV8-CTRL mice and AAV8-BMP4 mice. Error bars represent SEM.

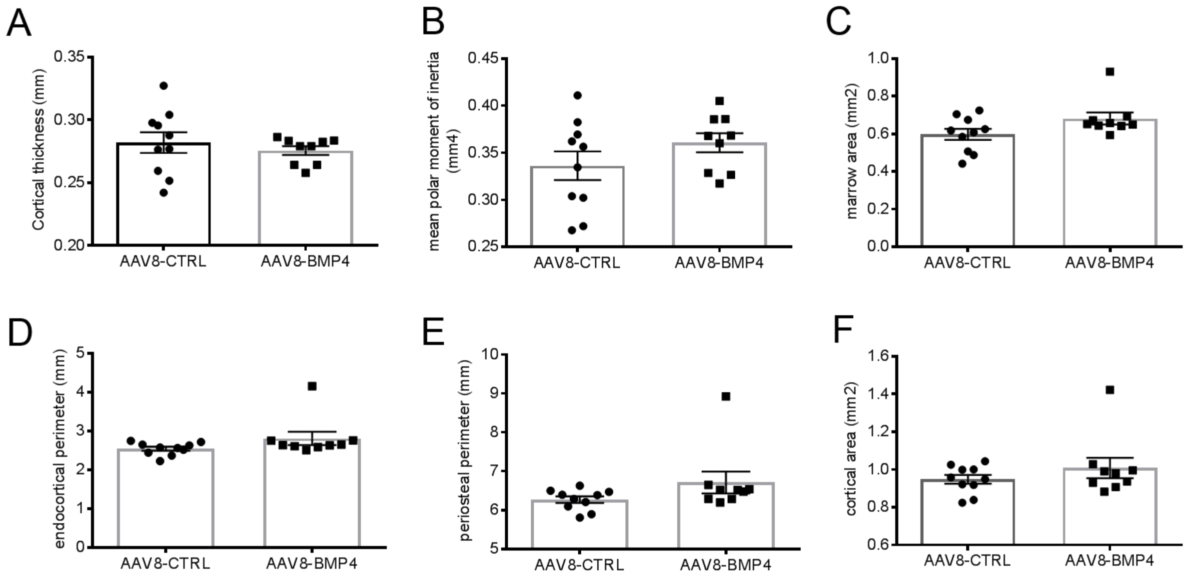


**Supplementary Figure 3. BMP4 effect on murine cortical bone.** The left femur from each mouse, AAV-CTRL (n=10) and AAV8-BMP4 (n=9) was harvested and examined *by ex vivo* µCT. (A) Cortical thickness, (B) mean polar moment of inertia, (C) marrow area, (D) endochondral perimeter, (E) periosteal perimeter and (F) cortical area. Error bars represent SEM. A two-tailed unpaired t-test was used to test significance.

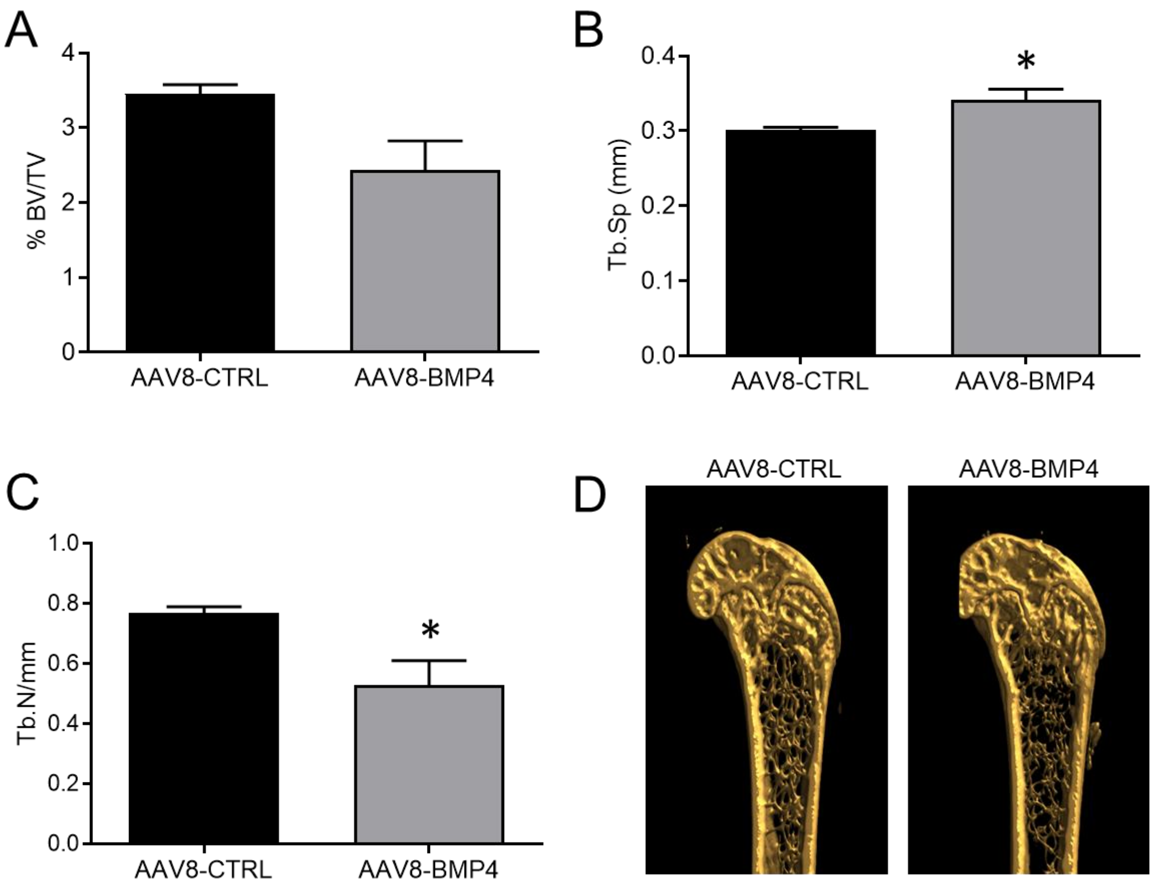


**Supplementary Figure 4. BMP4 effect on C57BL6/N murine bone.** C57BL6/N were injected with AAV8-CTRL or AAV8-BMP4 by tail-vein injections. At the end of the experiment one femur from each mouse, AAV8-CTRL (n=4) and AAV8-BMP4 (n=5), was harvested and examined by *ex vivo* µCT. (A) Trabecular volume as a proportion of tissue volume (BV/TV, %), (B) trabecular separation (Tb.Sp, mm) and (C) trabecular number (Tb. N, $mm^{-1}$) was assessed. Error bars represent SEM. (D) Representative images for an AAV8-CTRL mouse and an AAV8-BMP4 mouse are shown. Significance was tested using a two-tailed, unpaired t-test, *; $p<0.05$.

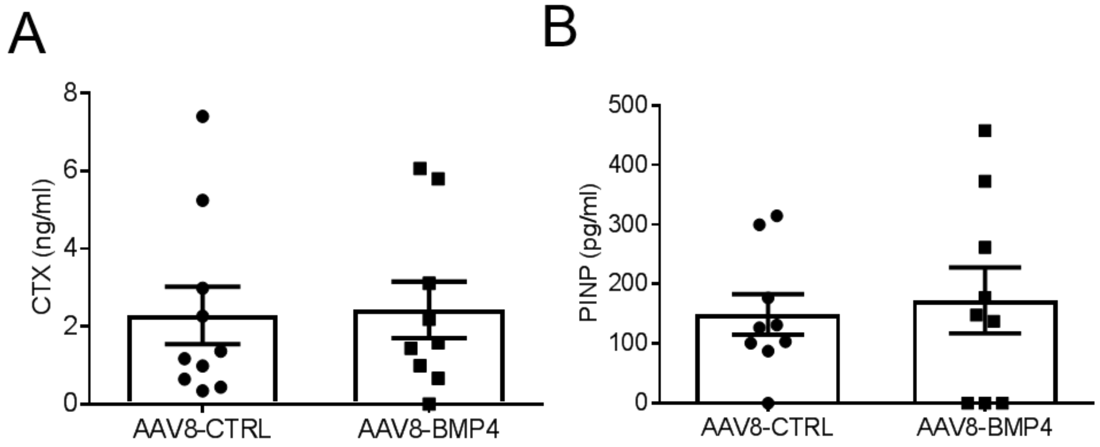


**Supplementary Figure 5. Serum levels of CTX-1 and PINP.** At end point, serum from AAV8-CTRL (n=10) and AAV8-BMP4 (n=9) was harvested and markers for (A) bone degradation (CTX-1) and (B) bone formation (PINP) was measured. Error bars represent SEM. Significance was tested using a two-tailed, unpaired t-test.

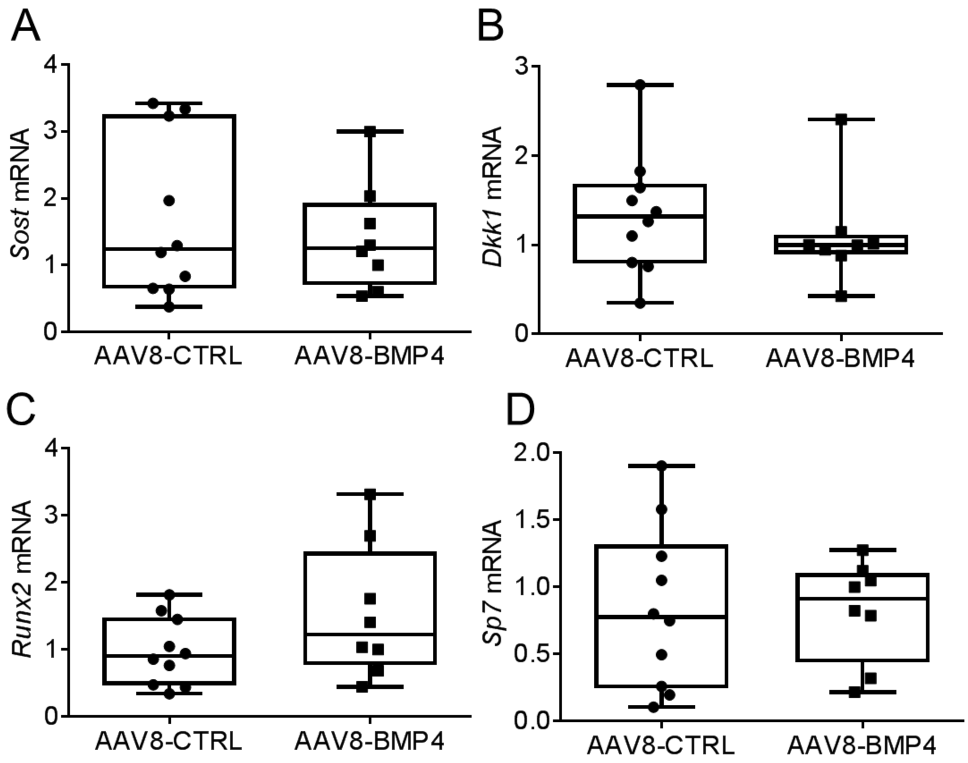


**Supplementary Figure 6. Expression of osteoblast specific markers in femurs.** Femur cDNA from AAV8-CTRL mice (n=10) and AAV8-BMP4 mice (n=8) was used for comparative RT-PCR using TaqMan Assays for the osteoblast specific markers *Sost* (A), *Dkk1* (B), *Runx2* (C), and *Sp7* (D). Gene expression was analyzed using the ΔΔCt method with *Gapdh* as housekeeping gene. There were no significant differences (two-tailed unpaired t-test).

**Supplementary Table 1. Taqman assays used for RT-PCR.**

| **Gene name** | **Assay #** |
|---|---|
| *Gapdh* | Mm99999915_g1 |
| *Smad7* | Mm00484742_m1 |
| *Id1* | Mm00775963_g1 |
| *Bmp4* | Mm00432087_m1 |
| *Bmp2* | Mm01340178_m1 |
| *Bmp6* | Mm01332882_m1 |
| *Gdf2* (BMP9) | Mm00807340_m1 |
| *Sost* | Mm00470479_m1 |
| *Dkk1* | Mm00438422_m1 |
| *Runx2* | Mm01269515_m1 |
| *Sp7* | Mm04209867_m1 |
| *Nfatc1* | Mm00479445_m1 |
| *Ctsk* | Mm00484039_m1 |
